# Charge density of cation determines inner versus outer shell coordination to phosphate in RNA

**DOI:** 10.1101/2020.03.10.986091

**Authors:** Hung T. Nguyen, D. Thirumalai

## Abstract

Divalent cations are often required to fold RNA, which is a highly charged polyanion. Condensation of ions, such as Mg^2+^ or Ca^2+^, in the vicinity of RNA renormalizes the effective charges on the phosphate groups, thus minimizing the intra RNA electrostatic repulsion. The prevailing view is that divalent ions bind diffusively in a non-specific manner. In sharp contrast, we arrive at the exact opposite conclusion using a theory for the interaction of ions with the phosphate groups using RISM theory in conjunction with simulations based on an accurate Three Interaction Site RNA model. The divalent ions bind in a nucleotide-specific manner using either the inner (partially dehydrated) or outer (fully hydrated) shell coordination. The high charge density Mg^2+^ ion has a preference to bind to the outer shell whereas the opposite is the case for Ca^2+^. Surprisingly, we find that bridging interactions, involving ions that are coordinated to two or more phosphate groups, play a crucial role in maintaining the integrity of the folded state. Their importance could become increasingly prominent as the size of the RNA increases. Because the modes of interaction of divalent ions with DNA are likely to be similar, we propose that specific inner and outer shell coordination could play a role in DNA condensation, and perhaps genome organization as well.

## 1 Introduction

How RNA molecules, containing only four bases with similar chemical properties, fold into a bewildering variety of complex structures continues to be a fascinating and unsolved problem.^1,2^ Because RNA molecules are polyanions, they require counter ions to fold. In particular, divalent cations are required to mute the repulsive electrostatic interactions between the negatively charged phosphate groups so that RNA can reach the folded state. As a consequence, ion–RNA interactions during the folding process has been the subject of extensive experimental and theoretical studies.^3–21^

From the theory of ion-induced shape changes in polyelectrolytes (PEs), it follows that divalent ions (such as Mg^2+^ and Ca^2+^) should be more efficient than monovalent ions in neutralizing the phosphate charges in RNAs.^22,23^ The physically appealing counterion condensation (CIC) theory could be used to estimate the fraction of condensed ions onto PEs with regular (rod-like or spherical) shape, and account for the phenomenon of entropically-driven counterion release in which monovalent ions are replaced by divalent ions. However, the CIC theory cannot be used in problems involving dramatic shape changes (RNA folding or PE collapse) as the ion concentration is altered. More importantly, the CIC assumes that the counterions uniformly decorate the polyanion, which is certainly valid for rod-like PEs. Because of the clear physics of the CIC theory, the prevalent view in the RNA literature is that most of the ions are uniformly (sometimes referred to as diffusive ions) condensed onto RNA. Although the CIC theory does estimate the extent of charge renormalization of the phosphate groups,^4,24^ it gives qualitatively incorrect picture of the mechanism by which divalent cations drive RNA folding.^2,25^

Recently, simulations of a large ribozyme and several other RNA constructs, in the presence of both monovalent and divalent ions, have shown that the ions have a strong propensity to bind to nucleotides in a site-specific manner, and not uniformly.^25–27^ The irregular shape of the RNA plays an important role in determining where the ions bind, as the ions are naturally drawn to the highly negatively charged pockets. Other studies, focusing only on the folded states or well-defined complexes involving RNA, have shown that the site-specifically bound Mg^2+^ ions to various RNA molecules are stable.^28–33^ These studies did not consider ion-driven folding of RNA from a completely unfolded state, which is the subject of our interest here and elsewhere.^25,26^ In order to develop a quantitative theory of RNA folding, effects of ions and the associated conformational changes they induce in RNA must be treated on equal footing.

Conceptually, specific association of divalent ions to RNAs could be categorized into two binding types: inner shell (or inner-sphere) binding where the divalent ion is partially dehydrated to interact directly with one (or more) atom of the RNA and outer shell (or outer-sphere) where it fully retains its first hydration shell to bind the RNA (illustrated in Fig. 1A).^9^ Inner shell binding of Mg^2+^ has been mostly detected in relatively large RNAs, where the ions are often localized at the deepest regions inside the RNAs.^34^ Characterizing which binding types the ions utilize to bind RNAs is difficult, and requires reliable computational methods because currently experimental techniques cannot easily distinguish between the two coordination modes.

**Figure 1:**
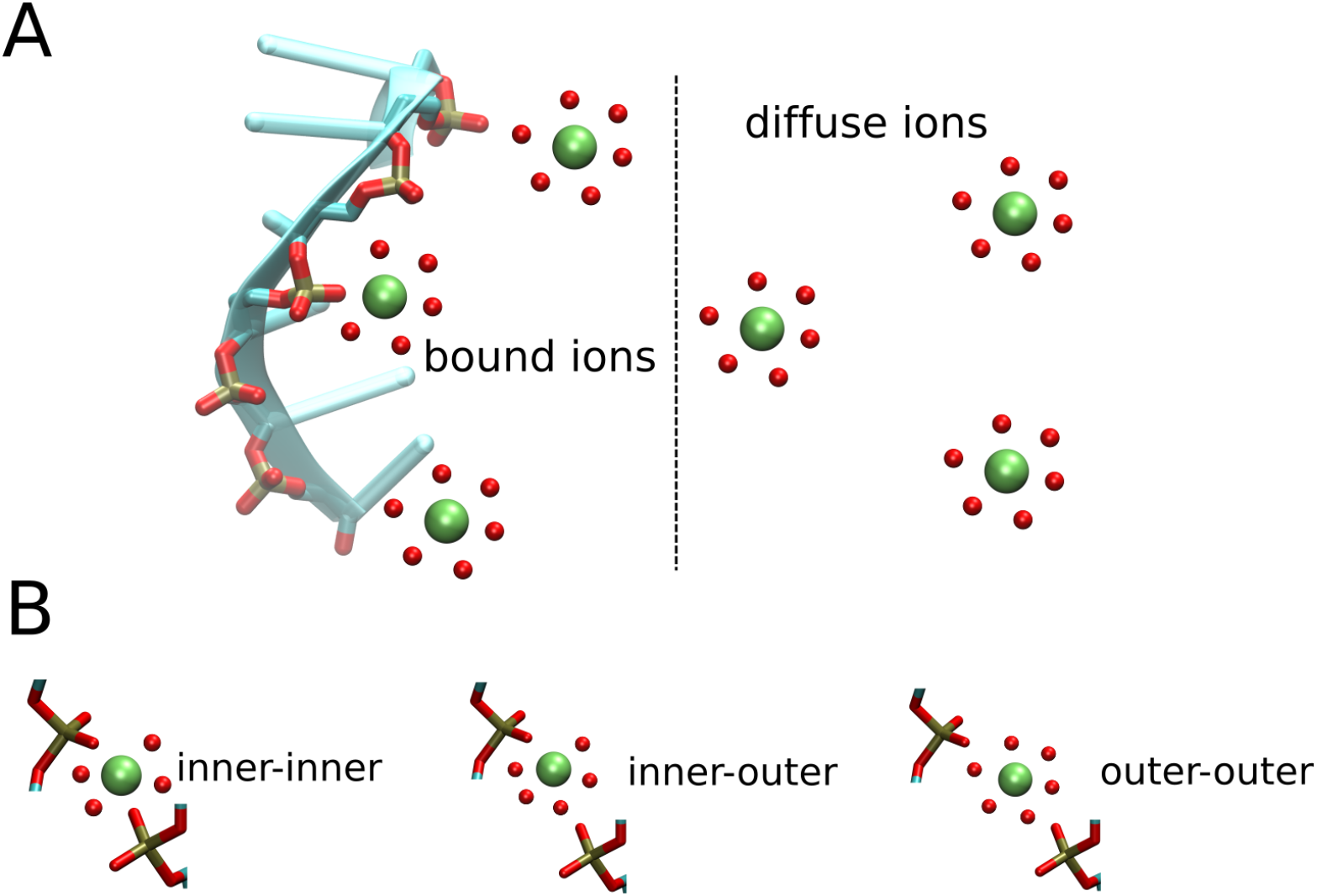
Schematic classification of divalent cations that interact with RNA. (A) Bound ions bind directly to the RNA in either the inner shell or outer shell binding. Diffuse ions, with intact hydration shells, are located further away from the RNA surface, interacting mainly with the RNA electrostatic field. For clarity, only water molecules in the first hydration shell of the ions are shown. (B) In some cases, a single ion could bind two phosphate groups simultaneously, resulting in bridging interaction. We refer these ions as bridging ions.

Divalent ion-driven folding of RNA is a strongly interacting many body problem because the binding of several Mg^2+^ ions in a specific and correlated manner is needed to obtain a stable folded structure.^25,26^ Already at the single ion level, the interplay between the inner and outer sphere coordination manifests itself. In a pioneering series of studies conducted nearly thirty years ago, Rossky and coworkers used extended reference interaction site model (RISM) theory and simulations to show that the potential of mean force (PMF) between Na^+^ and phosphate exhibits the two binding modes mentioned above.^35–37^ Building on these studies, we have recently developed a theory accounting for both types of binding between divalent ions and phosphate groups in RNAs.^38^ The theory, when incorporated in a thermodynamically accurate Three Interaction Site (TIS) force field for RNA,^39^ quantitatively reproduced ion-dependent RNA folding thermodynamics for several RNA molecules with different sizes, ranging from a small pseudoknot to the aptamer domain of the adenine riboswitch. We showed that both the inner and outer shell bindings are needed to produce a faithful description of ion-RNA interactions, because the ion-RNA interaction free energies, and therefore the RNA folding process depends sensitively on the ion-phosphate potential.

In this study, we focus on the mechanism of divalent cation condensation around RNA in order to in-vestigate the ion binding modes to the phosphate groups (inner or outer shell). We show that the divalent ion charge density as well as the RNA size determine the extent of inner versus outer shell binding to the phosphate groups. For small RNAs, such as the Beet Western Yellow Virus (BWYV) pseudoknot (PK), Mg^2+^ ions show a stronger propensity to bind to the outer shell. For larger RNAs, such as the 58-nucleotide fragment of the ribosomal RNA (58-nt rRNA), there is a shift in favor of the inner shell binding, because of an increase number of nearby phosphate groups in a larger RNA presumably compensating for the high penalty of ion dehydration. In contrast, Ca^2+^ ion, which has a larger radius and smaller charge density compared to Mg^2+^, dehydrates more readily to bind directly to the phosphate groups in the inner shell regardless of the RNA size. We also find that the bridging divalent ions, which bind two (or more) non-neighboring phosphate groups simultaneously, play a crucial role in stabilizing the RNA tertiary structure. Diluting the divalent ion concentration leads to a smaller probability of finding these bridging ions, thus destabilizing the folded state.

## 2 Materials and Methods

### 2.1 Divalent ion–phosphate interaction

Accurate simulations using coarse-grained models of ribozymes in explicit monovalent and divalent cations is computationally demanding.^25,26^ In order to simplify the problem, while still retaining a high level of accuracy in predicting the RNA folding thermodynamics and kinetics, we developed a theory to treat the electrostatic interactions involving monovalent ions implicitly, while treating the many body effects due to divalent ions explicitly.^38^ Briefly, we assumed that the screening effect due to monovalent cations, especially when present at high concentrations, can be described by the classical Debye–Huckel (DH) theory. However, an accurate theory is required to treat the short-ranged divalent ion–phosphate interaction, where the ion-ion correlation and charge polarization are important. We used RISM^35–37,40,41^ to calculate the short-ranged interaction between the divalent ion and the phosphate group, and smoothly join it to the Debye–Huckel potential at large separations. The resulting PMF between the cation X^2+^ and the phosphate group is given by,^38^

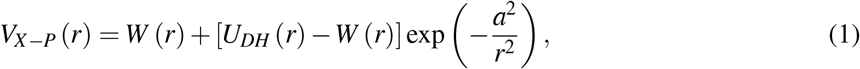

where *W* (*r*) is the PMF calculated using the RISM theory and *U*_*DH*_ (*r*) is the DH potential between X^2+^–P, accounting for the screening effect of monovalent ions. Our earlier work showed that the use of Eq. 1 in conjunction with the TIS model for RNA quantitatively reproduced the folding thermodynamics of several RNA molecules over a wide range of monovalent and divalent ion concentrations.^38^ Separation of long range potentials into a short-ranged part and the smoother long-ranged part had been previously proposed in several insightful studies.^42,43^ More recently, Gao *et. al.* have applied the Local Molecular Field treatment to calculate the PMF between Ca^2+^ and Cl^−^.^44^ The interesting finding is that a judiciously chosen short range potential in a system dominated by electrostatic interactions suffices to capture the physics of cation-anion interactions. Our work, which is similar in spirit, uses RISM to calculate the PMF between Mg^2+^/Ca^2+^ and phosphates in the monovalent salt buffer, which is needed to investigate RNA folding.

### 2.2 RNA force field and simulation details

#### Model

Since the details of the force field and simulations are given elsewhere,^38^ we only provide a brief description here. We used the TIS model in which each nucleotide is coarse-grained by three beads corresponding to phosphate, sugar and base moieties.^39^ The energy function in the TIS model is *U* = *U*_*BA*_ + *U*_*EV*_ + *U*_*ST*_ + *U*_*HB*_ + *U*_*EL*_, where *U*_*BA*_ takes into account the bond and angle restraints between connected beads, *U*_*EV*_ is the excluded volume term. Stacking and hydrogen bond interactions, *U*_*ST*_ and *U*_*HB*_, were calibrated to reproduce the heat capacity curves of small RNAs. ^45^ Once the two parameters were determined, we kept them unchanged for all RNA constructs. Thus, the model is transferable and could be used to study the folding of much larger RNA molecules.^25,26^ The electrostatic term *U*_*EL*_ is the sum of the Debye–Huckel energies for all the charges excluding the divalent ion–P term, which is calculated using Eq. 1.

#### Simulations

We performed simulations by integrating the Langevin equations of motion. Divalent cations were initially added randomly to a cubic box containing a RNA molecule of interest. The initial coordinates of the RNA were taken from the structure of the folded state in the PDB, 1L2X and 1HC8 for the 28-nt BWYV PK and 58-nt rRNA, respectively. The box size varied from 700-3,000 Å depending on the bulk concentration of divalent cations. We used large enough boxes to ensure that at least 200 divalent cations were present in the simulations. We used periodic boundary conditions to minimize the effect of finite box size. Numerical integration of the equations of motion was performed using the leap-frog algorithm. We carried out low-friction dynamics in order to increase the sampling efficiency of the conformations, in which the viscosity of water was reduced 100 times.^46^ Snapshots were taken every 10,000 steps, and the last two-thirds were used to calculate various quantities of interest.

#### Analyses

The local concentration of divalent ion around the i^th^ nucleotide in the RNA was computed using

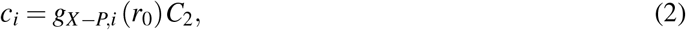

where *C*_2_ is the bulk concentration of divalent ion and *g*_*X−P,i*_ (*r*) is the radial distribution function of X^2+^ around the i^th^ phosphate group. The value *r*_0_ corresponds to the first or second peak in *g*_*X−P,i*_ (*r*), depending on the binding mode.

## 3 Results

### 3.1 Divalent cation distribution around RNA is site-specific

Fig. 2A shows the concentration profiles of Mg^2+^ from the center of BWYV PK. Although the bulk concentration is low (on the order of sub-mM), the local concentration of Mg^2+^ ions is ~ 3 orders of magnitude larger, indicating that the ions are strongly attracted to the PK. Interestingly, Figs. 2B and 2C illustrate that the distribution of Mg^2+^ ions around BWYV is not uniform, but site-specific. The fingerprints in Fig. 2B exhibit peaks at nucleotide positions G7, G12 and C22 for the folded state. These nucleotides are located at the central positions in the RNA structure (shown in Fig. 2C), where there are several phosphate groups surrounding them, thus resulting in strong negative electrostatic potential. These three residues glue the two stems and loops together, maintaining the structural stability of the PK. Had the ion condensation been uniform then the peaks in Fig. 2B would be absent, and the distribution would be uniform, which clearly is not the case.

**Figure 2:**
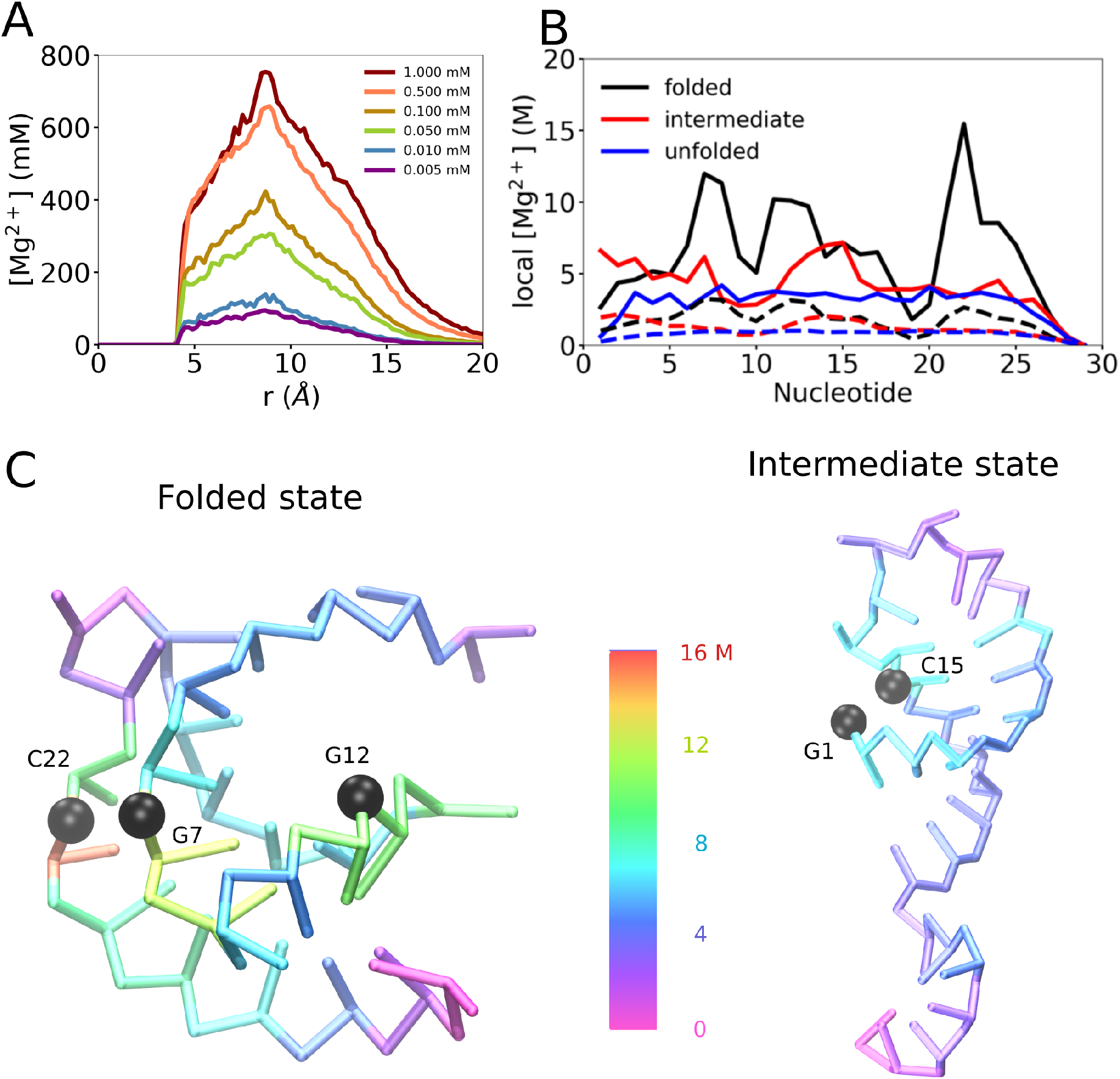
(A) Concentration profiles of Mg^2+^ relative to the geometric center of BWYV pseudoknot at various bulk concentrations. (B) Local concentration (Eq. 2) of Mg^2+^ in the inner shell (solid lines) and outer shell (dashed lines) computed for each nucleotide position at 54 mM KCl and 1.0 mM Mg^2+^. (C) Map of inner shell concentrations (M) onto representative snapshots of the folded and intermediate states at [Mg^2+^] = 1.0 mM. Phosphate groups of a few key nucleotides are shown in black spheres.

The Mg^2+^ concentration profiles for the intermediate and unfolded states are different compared to the folded state despite having the same bulk concentration. Because of the extended conformations, the unfolded state does not show any interesting feature in the profiles, and the Mg^2+^ ions have almost uniform binding affinities to all the nucleotides except near the 5’ and 3’ regions where the chain-end effects might be important. The profile of the intermediate state, adopting mostly hairpin conformations (shown in Fig. 2C), shows peaks at nucleotides G1 and C15, because of the formation of the hairpin leading to a high accumulation of phosphate groups around these nucleotides. The presence of ion peaks in the intermediate states shows that as soon as the titration starts, Mg^2+^ binds to specific nucleotides with high probability. When the RNA folds from the intermediate state to folded state, there is an interesting shift in the fingerprint of Mg^2+^.

Similar binding patterns are also observed for the case of 58-nt rRNA (Fig. 3). In this case, because the RNA folding depends on Mg^2+^ ions,^38,47^ new Mg^2+^ binding sites emerge as the ion concentration increases, leading to significant RNA compaction. Since the conformational changes occur when Mg^2+^ (or monovalent ion) concentration varies, one might expect there should be a binding pattern shift if Mg^2+^ concentration changes. Indeed, this is the case. Fig. 3A shows the local concentration of Mg^2+^ ions in the inner shell at two different bulk concentrations, 1.0 mM where the rRNA is fully folded and 0.02 mM where the rRNA mostly samples secondary structures. There are some notable differences in the two concentration profiles, especially at nucleotide positions that play key roles in maintaining the RNA tertiary structure. In the folded structure, G121 is located right at the intersection between the two stems and plays an extremely important role bridging the tertiary structure together, forming a Hoogsteen basepair with G141 and stacking on top of A139. (G137 glues the other two stems, but is located outside this region.) The phosphate groups of U147 and A148 point directly towards this region, where there is an overwhelming presence of other phosphate groups. This suggests that this region has a strong negative electrostatic potential, thus requiring excessive counterions to neutralize and stabilize the overall structure. We note that it is this region that gives rise to the difference in Mg^2+^ fingerprints when the Mg^2+^ concentration varies.

**Figure 3:**
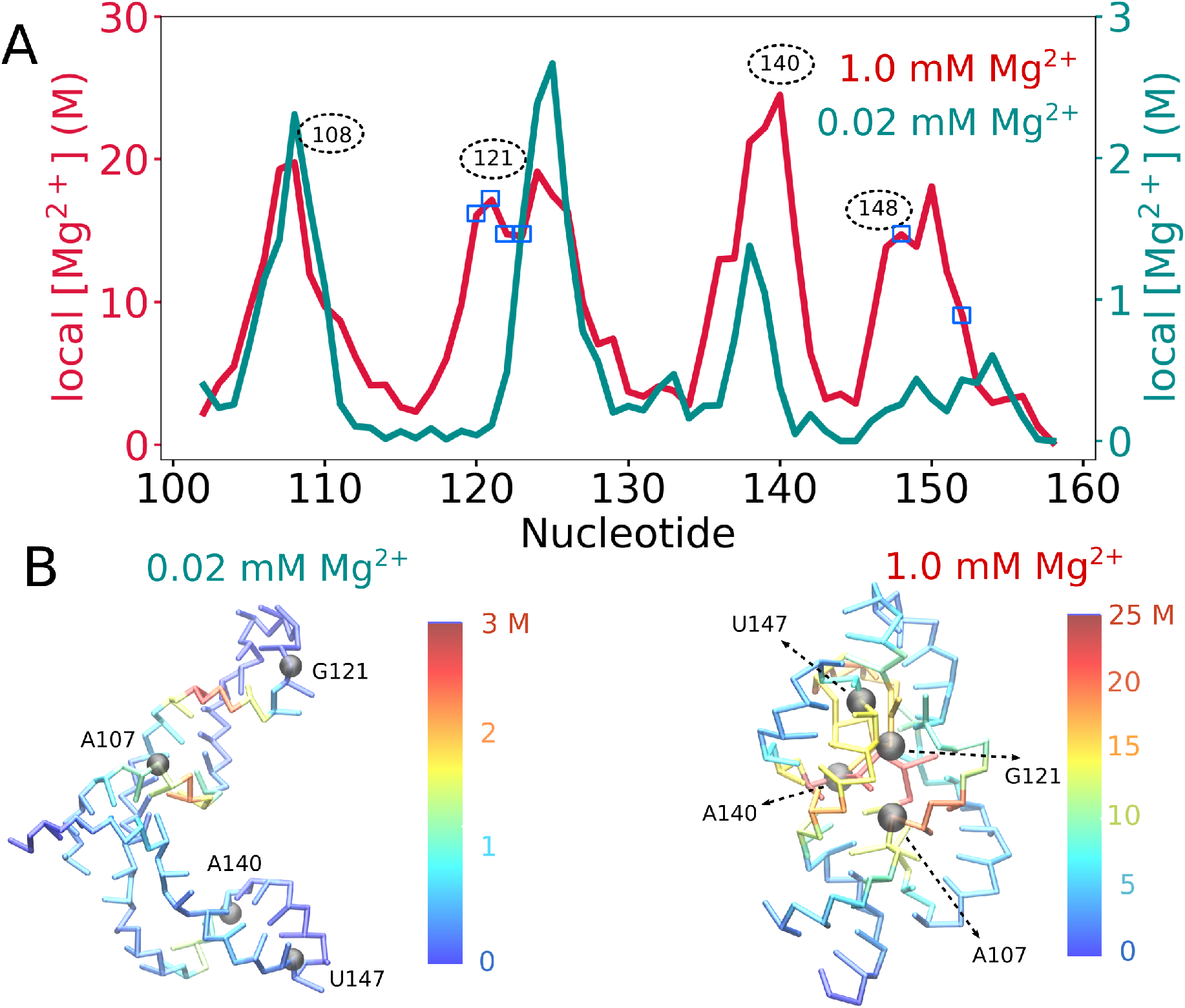
Fingerprint of Mg^2+^ binding to 58-nt rRNA. (A) Mg^2+^ local concentration in the inner shell at two concentrations 1.0 mM (red, left axis) and 0.02 mM (blue, right axis). Note that the scales associated with the two axes are different. Small blue squares are positions in the crystal structure where Mg^2+^ ions are found. At 1.0 mM Mg^2+^ concentration, the rRNA remains folded, whereas it partially unfolds at 0.02 mM Mg^2+^. Surprisingly, the ion binding patterns are similar except at some key nucleotides: A107, G121, A140 and U147, which are located at important positions in the rRNA structure. The phosphate groups in these nucleotides are highlighted as gray spheres. (B) Local concentration of Mg^2+^ ions mapped onto representative structures of the rRNA at 0.02 mM (left) and 1.0 mM (right) Mg^2+^ bulk concentration.

### 3.2 Inner shell *vs.* outer shell binding depends on the charge density of divalent ion

Do divalent ions interact with RNA directly or in the hydrated form? It is widely known that in aqueous solution, Mg^2+^ ion exists mainly in the hexahydrated form, Mg(H_2_O)_6_^2+^, because it has large favorable hydration free energy. Due to the high density charge, it would need a highly negatively charged environment (such as in large RNAs) in order to even partially dehydrate the Mg^2+^ ion. The lifetime of water coordinated to other divalent cations, which have a larger radius and therefore a smaller charge density, is relatively short. ^48^ Consequently, such ions (Ca^2+^ for example) readily dehydrate, and could bind directly to the phosphate groups through inner shell coordination. Currently, most (if not all) knowledge about the inner shell binding of divalent cations come mainly from X-ray crystallography, where the divalent ion location is assigned from the electron density map. Such an assignment for Mg^2+^ needs extra care to distinguish Mg^2+^ ion from isoelectronic species (having the same number of electrons) such as Na^+^, NH_4_^+^ and water, which are usually present in excess amount compared to Mg^2+^ in solution.^49,50^ Since our model describes both the inner and outer shell of Mg^2+^ around phosphate groups, we can determine how and where Mg^2+^ ions are localized around the phosphate groups in RNA.

Fig. 4A shows the radial distribution function between divalent ions and the P groups in BWYV PK. We performed the calculations for Mg^2+^ and Ca^2+^, although any spherical ion could be treated using our theory. It is obvious that our model accounts for both the inner and outer shell binding. We stress that it is crucial to treat both the binding modes on equal footing in order to reproduce ion–RNA binding thermodynamics.^38^

**Figure 4:**
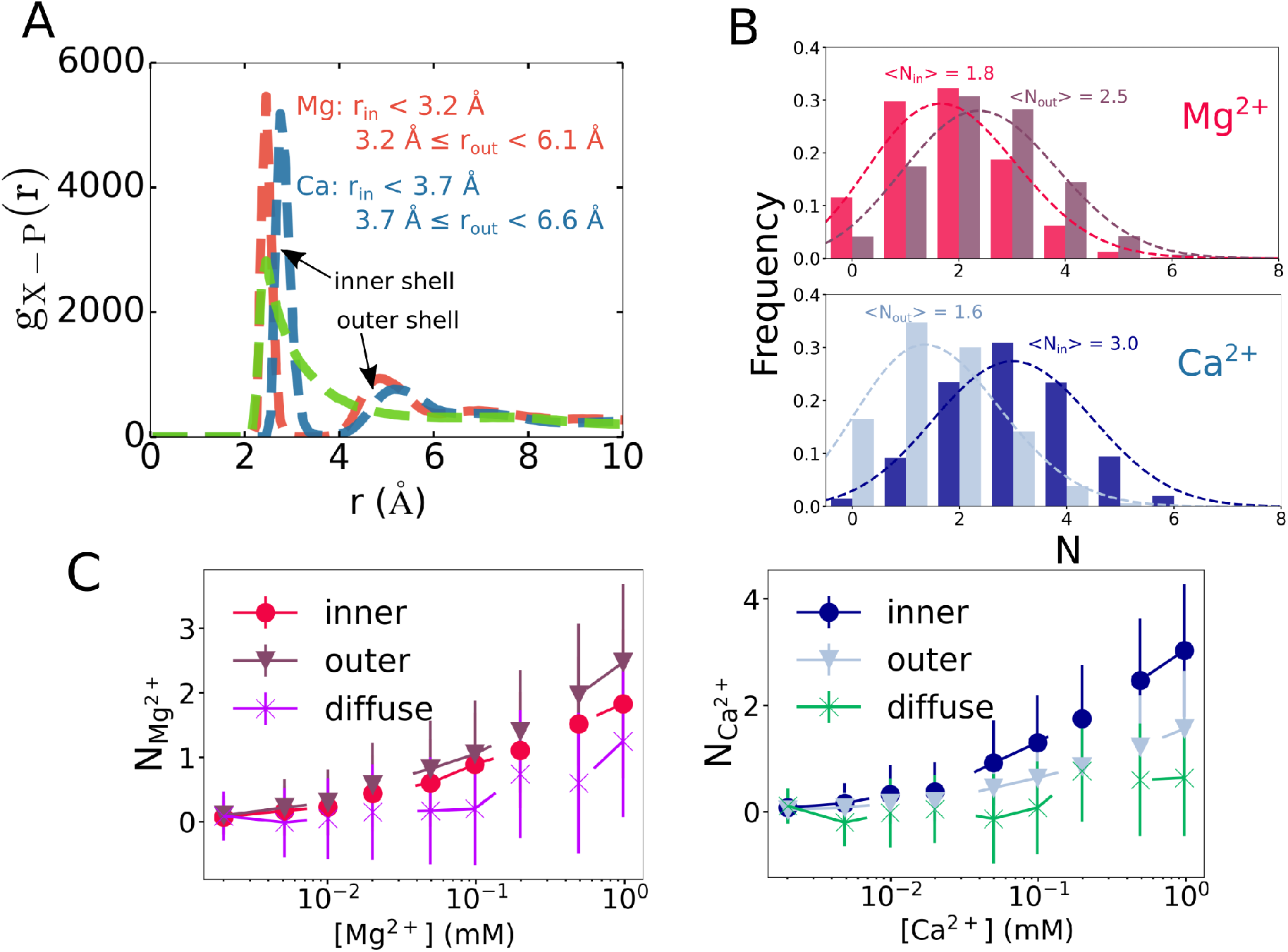
(A) Radial distribution function *g*_*X−P*_ (*r*) of divalent ions around phosphate groups in BWYV pseudoknot. Calculations were performed for 1.0 mM Mg^2+^ or Ca^2+^ in 54 mM KCl. For completeness, we also show *g* (*r*) for Mg^2+^ using the Debye–Huckel potential (green dashed curve). (B) Histogram of Mg^2+^ and Ca^2+^ ions in the inner, *N*_*in*_, and outer shell, *N*_*out*_. An ion is defined to be in inner or outer shell if the ion-phosphate distance falls within the first or second peak of *g*_*X−P*_ (*r*), respectively, as indicated in (A). (C) Averaged values of *N*_*in*_ and *N*_*out*_ at different bulk concentrations for Mg^2+^ and Ca^2+^. We also show the number of diffuse ions, which are located at least two hydration shells away from the RNA surface.

We computed the total number of ions in each binding mode for BWYV PK. If the ion-P distance falls within the first peak in *g*_*X−P*_ (*r*) (*r* < 3.2 Å for Mg^2+^, *r* < 3.7 Å for Ca^2+^) then the ion is considered to be in the inner shell. If it is within the second peak (3.2 *≤ r* < 6.0 Å for Mg^2+^, 3.7 *≤ r* < 6.6 Å for Ca^2+^) then the ion binds in the outer shell. Interestingly, some ions bind to the inner shell of one phosphate and to the outer shell of another phosphate group. We consider such ions as belonging to the inner shell. Fig. 4B shows that Mg^2+^ has a tendency to bind using the outer shell, even though we also find some ions that bind directly to the phosphates in the inner coordination. In contrast, Ca^2+^ prefers binding directly to phosphate groups without water-mediation. The higher charge density of Mg^2+^ ion, compared to Ca^2+^ ion, results in stronger interaction with water than Ca^2+^, thus shifting the binding pattern towards the outer shell interaction. The result is in agreement with the crystal structure of BWYV where there are totally 6 Mg^2+^ ions, and 3 of them bind directly to the phosphate groups, while the remainder 3 bind in the hexahydrated form. ^51^ It should be noted that, however, the conditions used in obtaining the crystal structure are different from those in the solution, and additional experimental data may be needed to confirm our predictions.

Fig. 4C shows the number of divalent ions found in the inner (*N*_*in*_) or outer shell (*N*_*out*_) at different bulk concentrations. We also calculated the number of diffuse ions, which are located at least two hydration shells away from the RNA surface, using *N*_*X,diff*_ = Γ_*X*_ −*N*_*X,in*_ −*N*_*X,out*_, where Γ_*X*_ is the preferential interaction coefficient for divalent ion X ^2+^, which is equivalent to the excess number of divalent ions attracted to the RNA, regardless of the position of the ions (data available elsewhere^38^). It can be seen from the results in Fig. 4C that the number of diffuse ions is low even at high bulk concentrations for both the ions, suggesting that divalent ions bind to RNA directly either via inner or outer shell binding. This agrees with our recent findings that in general, divalent cations are strongly attracted and bind RNAs directly and specifically, while the major role of monovalent ions is to screen the electrostatic propulsion between the phosphate groups.^25^ It also supports the picture from X-ray scattering measurements that divalent cations are generally localized closer to the RNA/DNA than monovalent ions.^52,53^

An interesting result is revealed when we investigate ion binding to the 58-nt rRNA, which is larger than BWYV. Fig. 5A shows that for this RNA, Mg^2+^ partially dehydrates and binds to the phosphates more readily using the inner coordination than the outer shell binding. The difference could be explained by the following arguments. BWYV is a relatively small 28-nucleotide RNA with simple open structure (Fig. 2C). The majority of the Mg^2+^ ions, consequently, stays near the RNA surface because there is no deep pocket with high electrostatic potential. As a result, only a couple of ions, which are located near the RNA center, dehydrate and bind using the inner shell. On the other hand, the rRNA is more complex, with a structure that has more deep pockets with highly negative electrostatic potential surrounded by a large number of phosphate groups (Fig. 3B). These regions need to be stabilized by small sized ions with large charge density in order for the RNA to stabilize the folded state. The high penalty associated with Mg^2+^ dehydration is partly (or even fully) compensated if the ion is attracted towards these regions. This could also partly explain why the 58-nt rRNA requires Mg^2+^ to fold. On the other hand, the majority of Ca^2+^ ions predominantly interacts directly with phosphate groups using the inner shell coordination. Compared with BWYV, the binding preference of both the ions features a slight shift towards the inner shell, which is expected for larger RNA molecules that require divalent ions to maintain their structural integrity.

**Figure 5:**
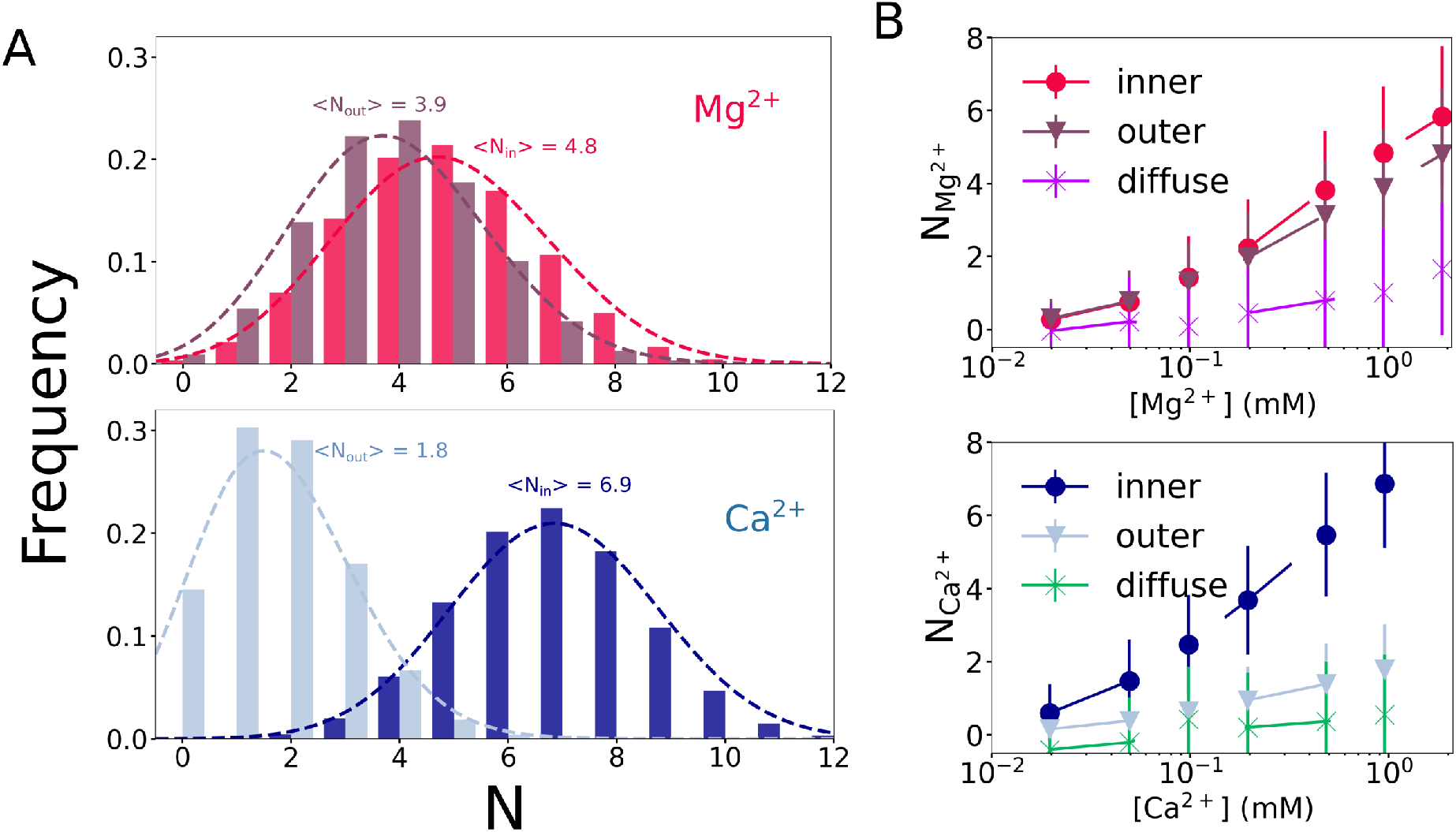
(A) Histograms of Mg^2+^ and Ca^2+^ coordination to the inner and outer shell at 1.0 mM divalent ions concentration for the 58-nt rRNA. Just as in the case for BWYV, Mg^2+^ binds to the rRNA using both inner and outer shell coordinations, whereas Ca^2+^ strongly favors the inner shell binding. (B) Averaged values of *N*_*in*_, *N*_*out*_ and the number of diffuse ions at different bulk concentrations for Mg^2+^ and Ca^2+^. Remarkably, the average number of diffuse ions is negligible, which further underscores the importance of specific binding.

### 3.3 Bridging divalent cations contribute to RNA stability

Ions that interact with RNAs are usually classified as diffuse and bound ions (Fig. 1). The bound ions can be further divided into outer shell and inner shell binding. Bridging interactions come from a bound ion that simultaneously interacts with two (or more) phosphate groups. Both phosphate groups could bind the ion in the inner shell or the outer shell, which we call inner-inner or outer-outer bridging, respectively. In addition, the ion could also bind one phosphate in the inner shell, while coordinating with another using the outer shell. We refer to this mode of bridging as inner-outer shell bridging. As we showed in previous sections, the divalent cations interact with RNAs via exclusively specific and direct bindings, either by inner shell or outer shell coordination. Consequently, we expect that the probability of divalent ions staying in the bridging position is high. Because the folding of the 58-nt RNA (and presumably other larger RNAs) depends strongly on Mg^2+^ ions, we propose that the bridging ions would be crucial in bridging distant nucleotides in order to form stable tertiary structures.

We calculated the probability of finding the bridging ions for any two nucleotides in the rRNA. Fig. 6 plots the results for inner-inner, inner-outer and outer-outer bridging as a function of Mg^2+^ concentrations. At high Mg^2+^ concentrations (1.0 mM and 0.1 mM), the rRNA remains mostly in the folded state, whereas at low Mg^2+^ concentrations (0.01 mM), it partially unfolds and populates extended states. Several interesting points can be made from the results in Fig. 6. First, as the ion concentration decreases, the probability of finding the bridging ions goes down as well. (We note that the color intensity for different ion concentrations is different, therefore we refer to the color bars for the actual values of the probabilities. We intentionally chose the color scales to demonstrate the clear shift in the bridging pattern at low ion concentration conditions.) This holds for all inner-inner, inner-outer and outer-outer bridging. The obvious reason is that as the ion concentration is lowered, the equilibrium is shifted to the unfolded states. Therefore, the attraction between the RNA and Mg^2+^ ions is weakened, leading to a smaller number of condensed ions. Second, it is rare to find a Mg^2+^ ion dehydrating two water molecules in the hydration shell simultaneously in order to bind two phosphate groups in the inner shell. The largest probability we observed in the simulations for such an event is ~ 2% at the highest Mg^2+^ concentration (1.0 mM). More crucially, only Mg^2+^ ions residing near the core of the rRNA, where there is an overwhelming number of phosphate groups, are able to participate in inner-inner bridging (yellow and purple regions in Fig. 6). Such bridging is also seen in inner-outer and outer-outer bridging as well, albeit with a much higher probability, ~ 15%. Interestingly, as the Mg^2+^ concentration decreases, the probability of finding the bridging Mg^2+^ ions in this region decreases, leading to a destabilized RNA and unfolding occurs. Our simulations, therefore, suggest that the presence of excess Mg^2+^ ions in this region, especially bridging Mg^2+^, play a vital role in driving the RNA to the folded structure.

**Figure 6:**
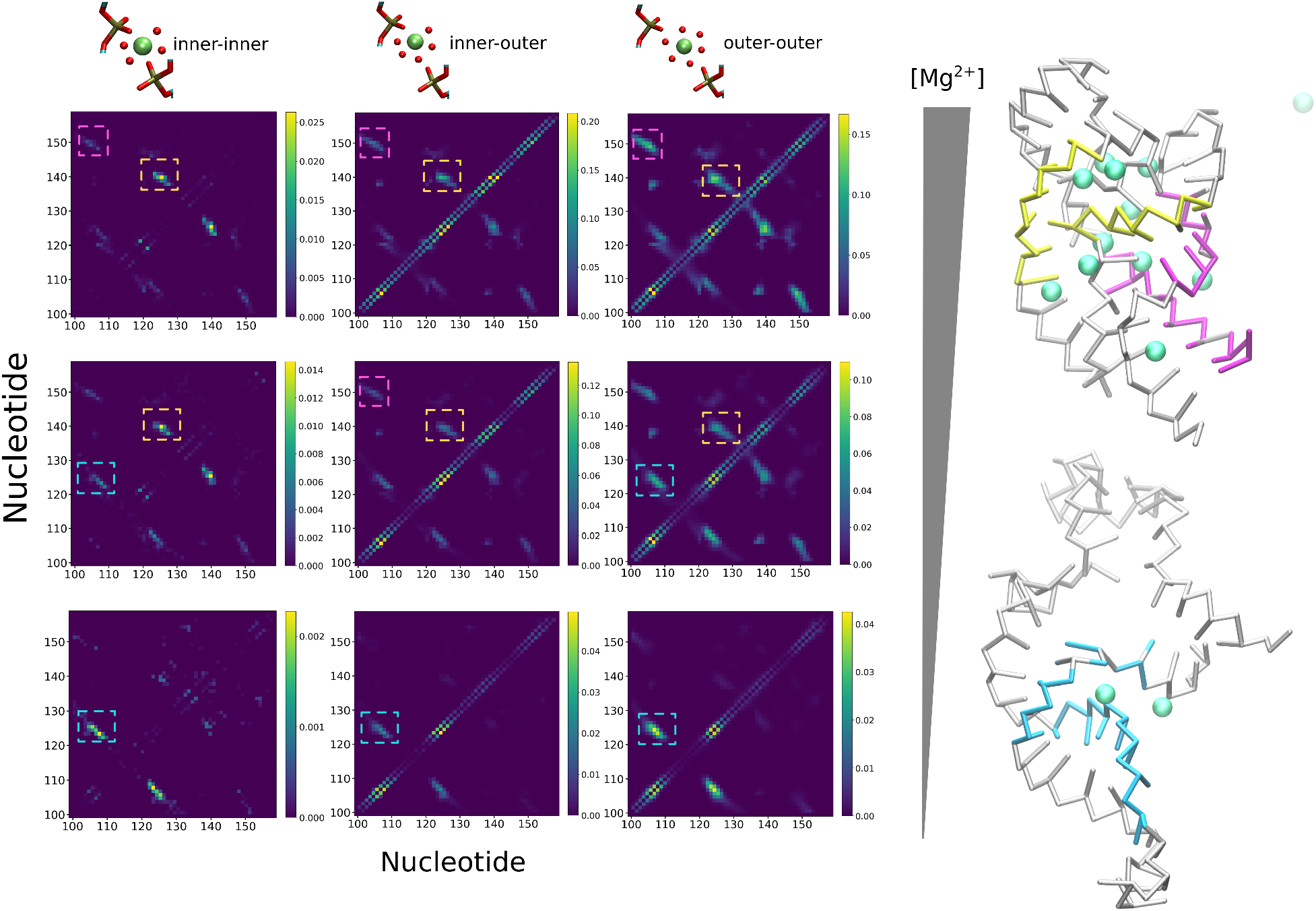
Role of bridging Mg^2+^ ions on 58-nt rRNA folding. Shown here is the heat map of where the bridging ions are found. The color bars on the right of each panel denote the bridging probability. On the left is the inner-inner bridging ion, where Mg^2+^ binds two phosphate groups in the inner shell simultaneously. Inner-outer bridging is shown in the middle, where Mg^2+^ binds one phosphate group in the inner shell and another using the outer shell coordination. In the outer-outer bridging (right), Mg^2+^ ions are coordinated to both groups using the outer shell. From top to bottom, the bulk concentration of Mg^2+^ decreases from 1.0 mM, 0.10 mM to 0.010 mM in a buffer containing 60 mM KCl. The color intensity is varied for different bulk concentrations, so the lower concentrations appears to have the same color intensity, but the actual probability is lower (see the color bars). Such a representation highlights the change in the bridging pattern as the RNA unfolds. The two most likely conformations on the right are in high and low Mg^2+^ concentrations, respectively. The Mg^2+^ ions are depicted as green spheres. Parts of the structures that are colored purple, yellow and blue are regions that form bridging interactions and are denoted by colored dashed squares in the heat map.

**Figure 7:**
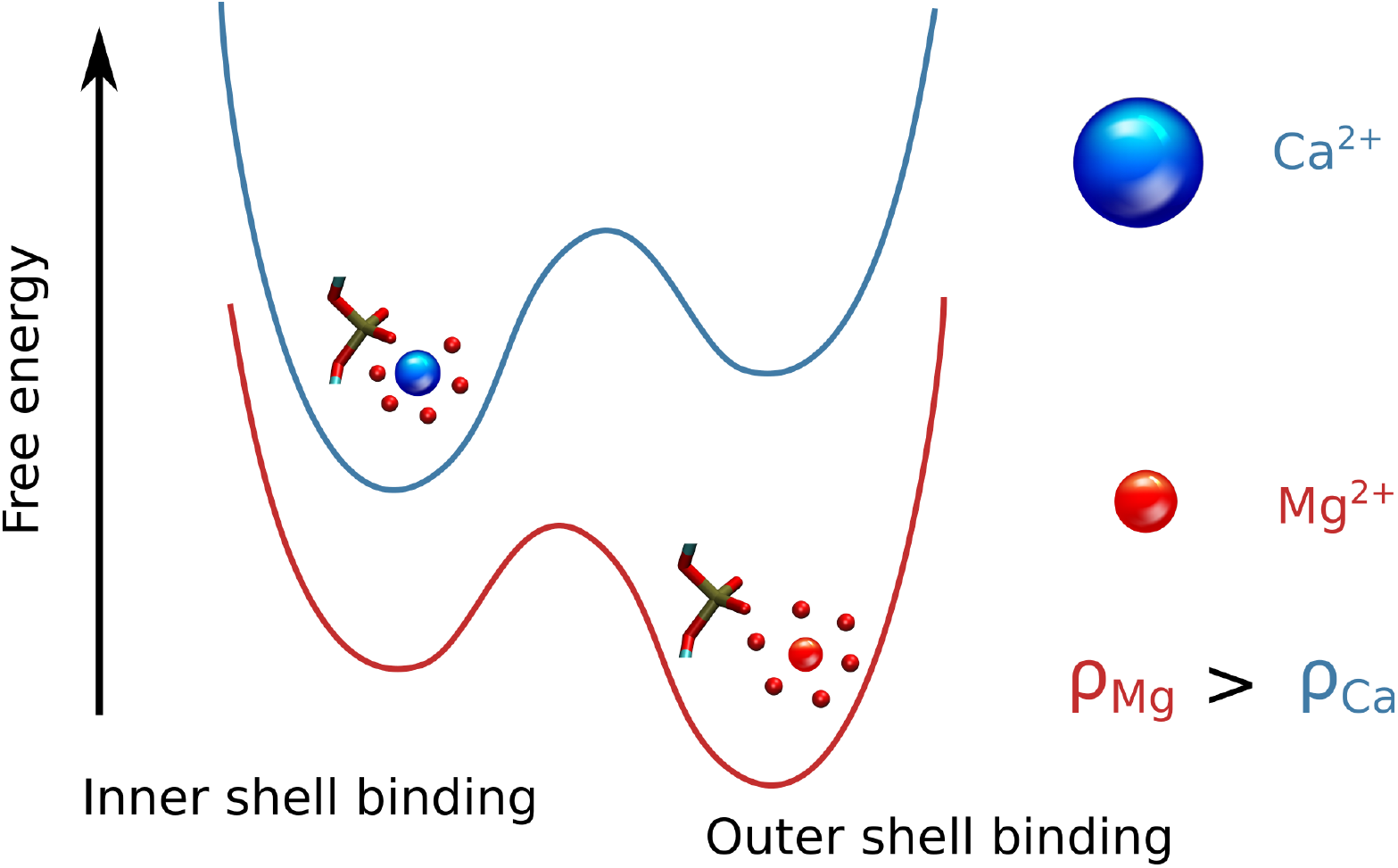
TOC image

## 4 Discussions and Conclusions

### Binding of Mg2+ and Ca2+ is nucleotide specific

Using simulations based on the accurate TIS model, which treats divalent cations explicitly and monovalent cations implicitly, we have established that divalent cations bind with high degree of specificity to the phosphate groups in RNA. This finding is in sharp contrast with the usual assumption that ions (often referred to as diffuse ions) are randomly localized around RNA merely to renormalize the charge on the phosphate groups. This assumption is based on the CIC theory that is strictly applicable to only rod-like or spherical polyelectrolytes (PEs),^22,23^ and not to PEs with irregular shapes containing groves. In the context of RNA folding, we find that the binding specificity depends on the nucleotide position in the folded state of the RNA. Surprisingly, the specificity of binding also depends on the nature of the intermediate states populated along the folding pathways. This implies that even at extremely low divalent ion concentrations, interactions with RNA results in structure formation.^26^ More importantly, there is a high degree of ion-ion coordination in the association of divalent cations, which again depends on the architecture of the folded states.

We also find that the majority of divalent cations binds directly to the phosphate groups in RNA using inner or outer sphere coordination. The needed screening of phosphate charges is left to monovalent ions, which are always present in the buffer or can also be increased externally. This new physical picture, based on theory and simulations, explains why only a “trace” amount of divalent ions is needed to significantly enhance RNA stability compared to monovalent ions. A rule of thumb in several experiments is that roughly 100-unit of monovalent ions is needed to replace 1-unit of divalent ions. ^54^ For example, *Tetrahymena* ribozyme folds at around 1 M monovalent ion (perhaps not to the functionally competent state) but requires only ~ 5 mM divalent cation.^4,55^

### Divalent charge density dictates inner *vs.* outer shell binding

Perhaps, the most significant finding of the present study is that there are distinct chelation modes in the interaction of divalent cations with RNA during the folding process. To the best of our knowledge, the present work and a related previous study^38^ are the only ones to establish that RNA stability depends on the inner and outer shell binding of divalent ions to the RNA phosphate groups. In particular, we have shown that the charge density of the ions has a strong effect on how divalent ions bind to the phosphate groups in RNA. In BWYV PK, Mg^2+^ preferentially binds to the outer shell. However, with reduced probability, some Mg^2+^ ions are also localized near the RNA core, which requires modest dehydration to bind to phosphates in the inner shell. For the larger (58-nt) fragment of the rRNA, the RNA-generated negative electrostatic potential is high enough to compensate for the high dehydration penalty of Mg^2+^, resulting in more ions that bind to the inner shell. Indeed, in the 58-nt rRNA, the preferred mode of binding is to the inner shell. For more complex RNAs, it is likely that there would be even more Mg^2+^ ions binding directly to the phosphate groups. This accords well with the observation that most of the inner shell binding of Mg^2+^ ions is detected only for sufficiently large RNA structures.^34^

For Ca^2+^, which has a lower charge density compared to Mg^2+^, the situation is different. In particular, Ca^2+^ ions prefer binding directly to RNA using the inner shell coordination *regardless* of the size of the RNA molecules. Dehydration of Ca^2+^ ions is readily achieved because of the lower charge density, which makes it possible to coordinate with the phosphate group through the inner shell. The dramatic shift in the binding pattern illustrated here between the two divalent ions demonstrates the exquisite dependence of the ion-RNA interaction on the cation charge density, which has been also illustrated previously using simulations and experiment.^55^

### Bridging interactions play an important role in RNA stability

Divalent ions are required to bring non-neighboring phosphate groups together to facilitate tertiary structure formation. Our simulations show that this is indeed the case, and in the process the bridging ions play a crucial role in facilitating RNA folding. Once the divalent ion concentration is sufficiently low, leading to decreased probability of forming bridging ions, the tertiary structure loses stability, and the RNA eventually unfolds. Interestingly, the majority of Mg^2+^ ions engages in outer shell coordination if they bridge two non-neighboring phosphate groups. This finding is reasonable because it is would be free energetically prohibitive for Mg^2+^ to dehydrate twice in order to accommodate two phosphate groups in the inner shell. The only instance this occurs in our simulations is near the core of the RNA where there is limited space and an overwhelming number of phosphate groups. Even in this extreme case, the probability of the inner-inner bridging is very small, around 2%.

### Ion–DNA interactions

Toroid formation in DNA or, in general, DNA condensation (referred to as *Ψ* condensation) could also be driven by Mg^2+^.^56,57^ Because the theory presented here is general, we believe that Mg^2+^ coordination to the phosphate groups in DNA is likely to occur preferentially by outer shell binding. It would be most interesting to examine the sequence-specific binding of divalent cations to DNA in order to provide a molecular basis of Mg^2+^-induced flexibility and the mechanism of *Ψ* condensation. ^58,59^

## 5 Acknowledgments

It is a pleasure to dedicate this article to Peter Rossky whose profound contributions to our understanding of charged systems and many other topics have inspired one of us (DT) for decades. We are grateful to Naoto Hori for useful discussions. This work was supported by NSF Grant CHE 19-00093 and the Welch Foundation Grant F-0019 through the Collie–Welch chair. We thank the Texas Advanced Computing Center for providing computational resources.

